# Multi-pronged human protein mimicry by SARS-CoV-2 reveals bifurcating potential for MHC detection and immune evasion

**DOI:** 10.1101/2020.06.19.161620

**Authors:** AJ Venkatakrishnan, Nikhil Kayal, Praveen Anand, Andrew D. Badley, George M. Church, Venky Soundararajan

## Abstract

The hand of molecular mimicry in shaping SARS-CoV-2 evolution and immune evasion remains to be deciphered. We identify 33 distinct 8-mer/9-mer peptides that are identical between SARS-CoV-2 and human proteomes, along similar extents of viral mimicry observed in other viruses. Interestingly, 20 novel peptides have not been observed in any previous human coronavirus (HCoV) strains. Four of the total mimicked 8-mers/9-mers map onto HLA-B*40:01, HLA-B*40:02, and HLA-B*35:01 binding peptides from human PAM, ANXA7, PGD, and ALOX5AP proteins. This mimicry of multiple human proteins by SARS-CoV-2 is made salient by the targeted genes being focally expressed in arteries, lungs, esophagus, pancreas, and macrophages. Further, HLA-A*03 restricted 8-mer peptides are shared broadly by human and coronaviridae helicases with primary expression of the mimicked human proteins in the neurons and immune cells. This study presents the first comprehensive scan of peptide mimicry by SARS-CoV-2 of the human proteome and motivates follow-up research into its immunological consequences.

## Introduction

Viral infection typically leads to T cell stimulation in the host, and autoimmune response associated with viral infection has been observed^1^. SARS-CoV-2, the causative agent of the ongoing COVID-19 pandemic, has complex manifestations ranging from mild symptoms like loss of sense of smell (anosmia)^2^ to severe and critical illness^3,4^. While some molecular factors governing SARS-CoV-2 infection of lung tissues, such as the ACE2 receptor expressing cells have been characterized recently^5^, the mechanistic rationale underlying immune evasion and multi-system inflammation (Kawasaki-like disease) remains poorly understood^6,7^.

The SARS-CoV-2 genome encodes 14 structural proteins (e.g. Spike protein) and non-structural proteins (e.g. RNA-dependent RNA polymerase), as depicted in **Figure 1a**. The non-structural ORF1ab polyprotein undergoes proteolytic processing to give rise to 15 different proteins (NSP1, NSP2, PL-PRO, NSP4, 3CL-PRO, NSP6, NSP7, NSP8, NSP9, NSP10, RdRp, Hel, ExoN, NendoU and 2’-O-MT). The human reference proteome consists of 20,350 proteins, which when alternatively spliced, result in over 100,000 protein variants (**Figure 1b**)^8^.

**Figure 1.**
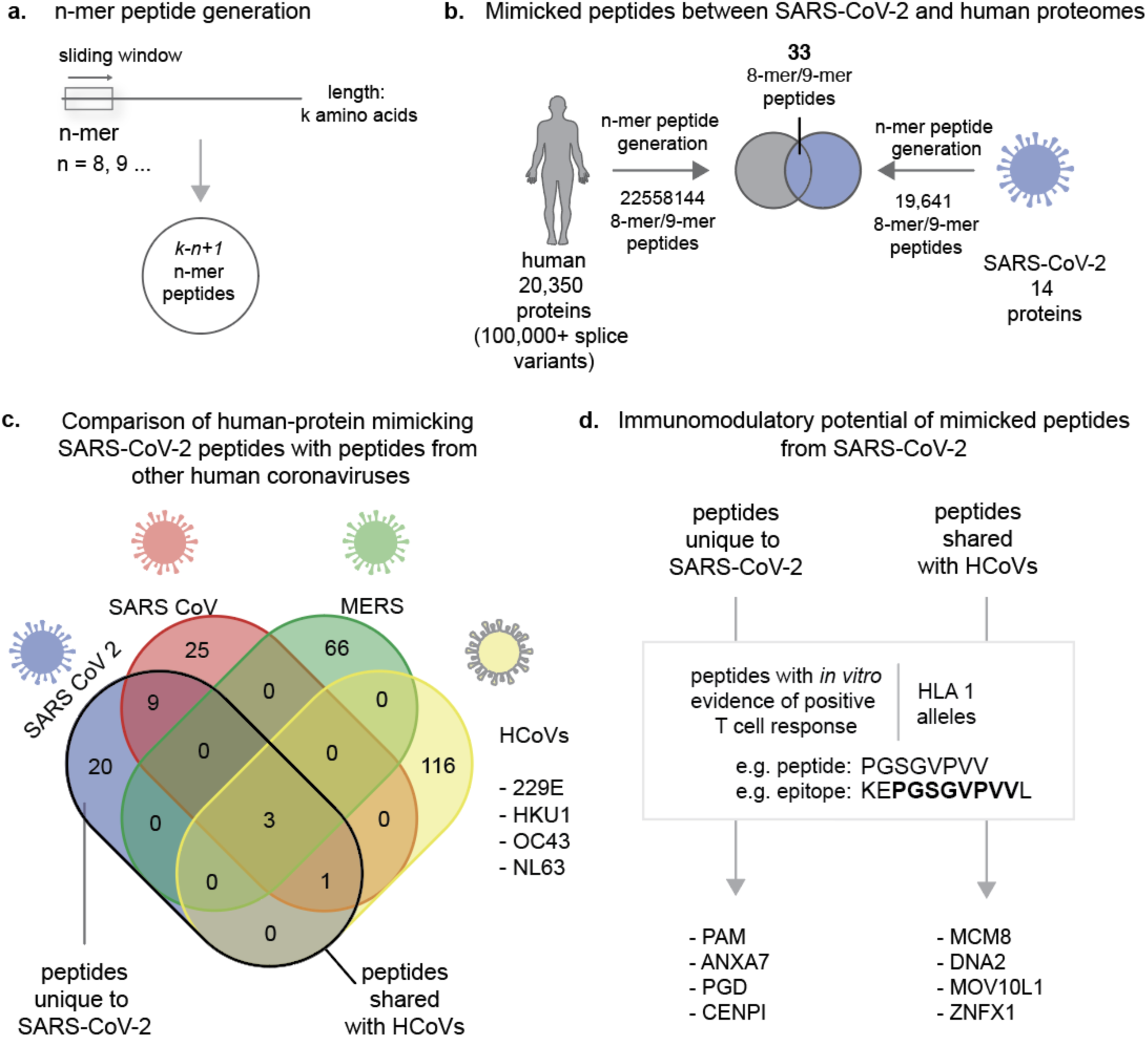
Molecular mimicry and immunomodulatory potential. (a) n-mer peptide generation. (b) Mimicked peptides between SARS-CoV-2 and human proteomes. (c) Comparison of human-protein mimicking SARS-CoV-2 peptides with peptides from other human coronaviruses (d) Immunomodulatory potential of mimicked peptides from SARS-CoV-2.

Here, we investigate the potential for molecular mimicry-mediated escape from immune surveillance and host antigen recognition in COVID-19, by performing a systematic comparison of known MHC-binding peptides from humans and mapping them onto SARS-CoV-2 derived peptides (see *Methods*). For this, we compute the longest peptides that are identical between SARS-CoV-2 reference proteins and human reference proteins, thus creating a map of COVID-19 host-pathogen molecular mimicry. Based on this resource, we extrapolate the potential for HLA Class-I restricted, T-cell immune stimulation via synthesizing established experimental evidence around each of the mimicked peptides.

## Methods

### Computing the SARS-CoV-2 peptides that mimic MHC Class-I binding peptides on the human reference proteome

The reference proteome for SARS-CoV-2 consists of 14,221 8-mers and 14,207 9-mers, that result in 9,827 distinct 8-mers and 9,814 distinct 9-mers (see **Table S1**). The 8-mers and 9-mers are generated by using a sliding window, moving one amino acid at a time, resulting in overlapping linear peptides. The reference human proteome, on the other hand, consists of 11,211,806 peptides that are 8-mers and 11,191,459 peptides that are 9-mers, resulting in 10,275,893 unique 8-mers and 10,378,753 unique 9-mers. Including alternatively spliced variants, increases the unique peptide counts to 11,215,743 8-mers and 11,342,401 9-mers.

Thirty one 8-mer peptides and two 9-mer peptides are identical between the reference proteomes of SARS-CoV-2 and humans, after including alternatively spliced variants (**Figure 1b, Table S2**). For comparison, we analyzed the protein sequences from 9434 viral species (taxons) from NCBI RefSeq (see *Methods*). On average around 45.57 unique 8mer/9mers per viral taxon were shared with the human proteome (mean = 45.57, median = 12, S.D. = ±135.38). In order to control for the complexity and constraints of the amino acid sequences, we also analyzed the distribution of mimicked 8-mers/9-mers normalized by the total number of unique 8-mers/9-mers present in each viral taxon. On average, a fraction of 0.002 8-mers/9-mers out of all the unique 8-mers/9-mers in the virus are identical between the viral proteome and the human proteome (mean = 0.002, S.D. =± 0.011)(**Figure 1 - Supplemental Figure 1**). The fraction of 33 human-mimicking 8-mers/9-mers is proximal to the mean. Overall, this suggests that the presence of 33 human-mimicking 8-mers/9-mers in SARS-CoV-2 is not surprising compared to the number of human-mimicking 8-mers/9-mers present in other viruses.

In SARS-CoV-2, no 10-mer or longer peptides are identical between the pathogen and the host reference proteins. Of these, 29 peptides (8-mer/9-mer) mimicked by SARS-CoV-2 map onto nine of 14 SARS-CoV-2 proteins and 29 of 20,350 human proteins. By including alternative splicing, the 33 8-mers/9-mers mimicked by SARS-CoV-2 map onto 39 of 100,566 protein splicing variants. That is, 0.16% of human proteins and 0.04% of all splicing variants have 8-mer/9-mer peptides that are mimicked by the SARS-CoV-2 reference proteome. Given that MHC Class I alleles typically engage peptides that are 8-12mers^9^, the analysis that follows was restricted to mapping host-pathogen mimicry from an immunologic perspective across 8-mer and 9-mer peptides only.

### Comparative analysis of SARS-CoV-2 peptides mimicking the human proteome with the reference SARS, MERS and seasonal HCOVs

Of the 33 peptides from SARS-CoV-2 that mimic the human reference proteome, 20 peptides are not found in any previous human-infecting coronavirus (SARS, MERS or seasonal HCoVs) (**Table 1**). The UniProt database was used to download the 15 protein reference sequences for SARS-CoV. The non-redundant set of protein sequences from other coronavirus strains (HCoV-HKU1:188; HCoV-229E: 246; HCoV-NL63: 330; HCoV-OC43: 910 and MERS: 681) was computed by removing 100% identical sequences, and the remnant sequences were all included in the comparative analysis with SARS-CoV-2 mimicked peptides. A Venn diagram depicting the overlap of mimicked peptides and identifying unique SARS-CoV-2 mimicked peptides was generated (**Figure 1c)**.

**Table 1.**
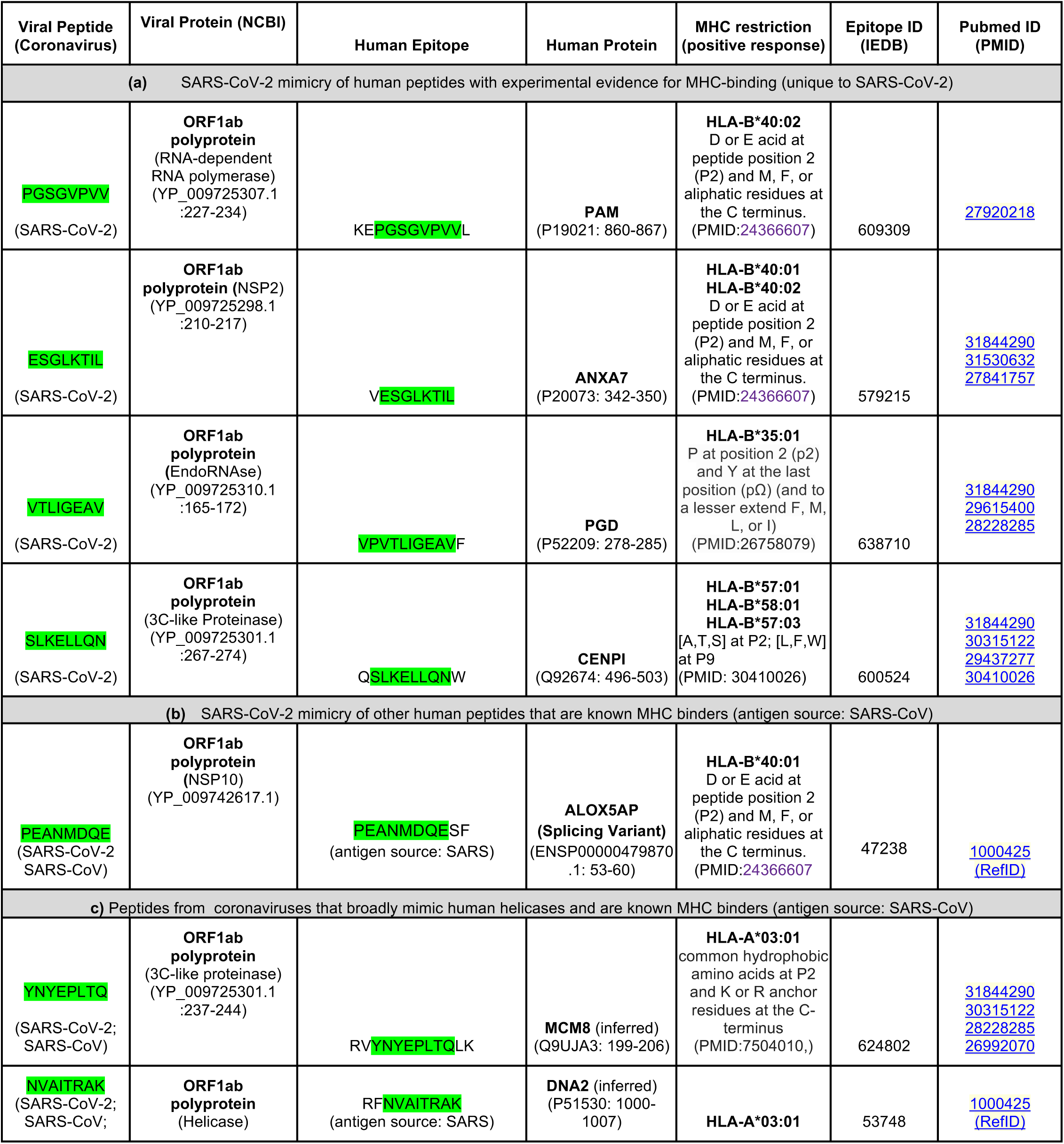
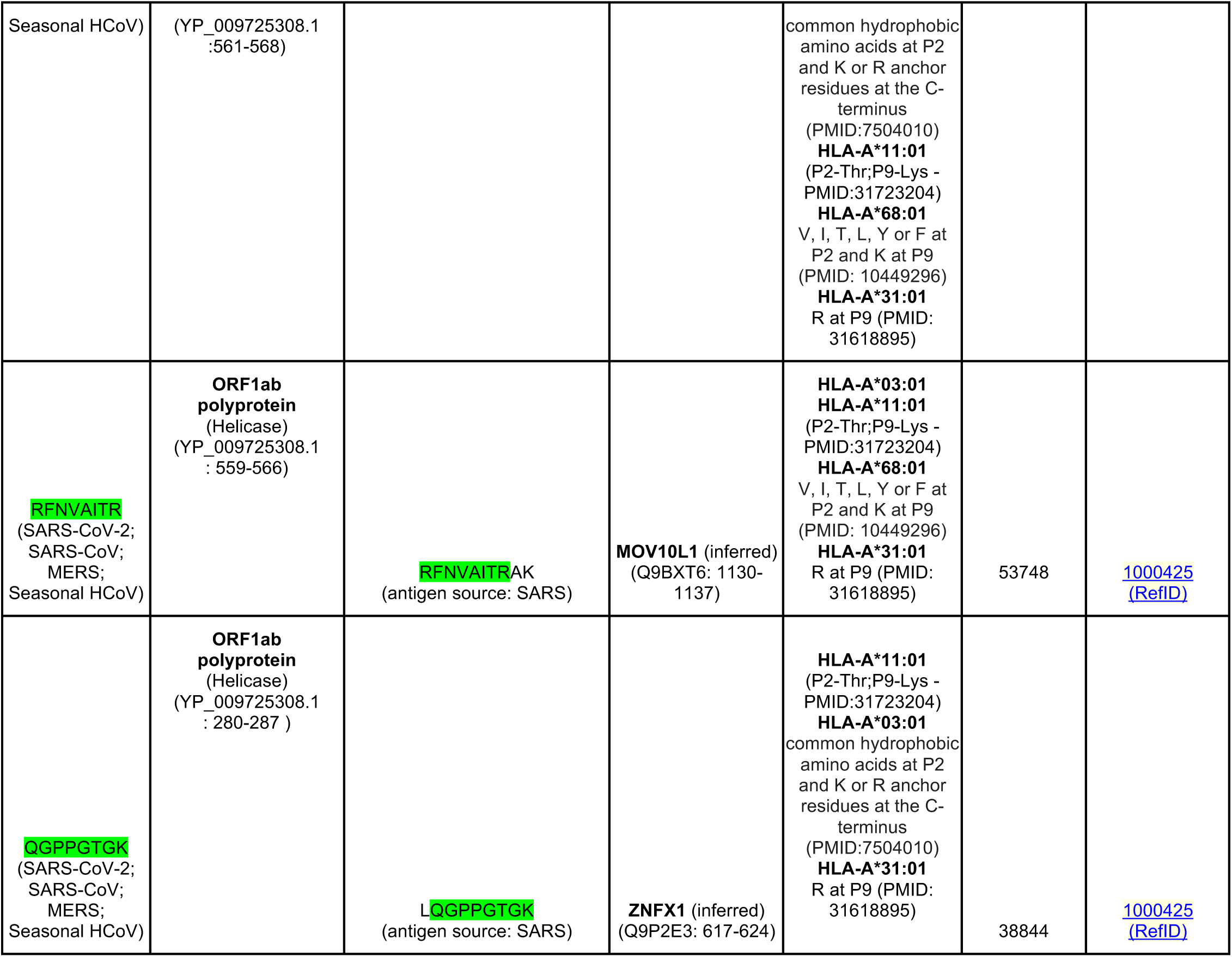
SARS-CoV-2 peptides mimicking human proteins, with experimental evidence of positive MHC binding from the immune epitope database. The viral-human mimicked 8-mer/9-mer peptides are highlighted in green text.

Given the zoonotic transmission potential of coronaviruses from other organisms to humans, we also considered 8-mer/9-mer peptides derived from 13,431 distinct protein sequences of all non-human coronaviridae from the VIPERdb^10^.

### Characterizing the SARS-CoV-2-derived 8-mers/9-mers that mimic established human MHC-binding peptides

The lengths of the human proteins mimicked by SARS-CoV-2 were considered to examine any potential bias towards the larger human proteins (**Figure 1-Supplementary Figure 1**). For instance, human Titin (TTN), despite containing 34,350 amino acids and being of the longest human proteins, does not have even one peptide mimicked by SARS-CoV-2. The longest and shortest human proteins that are mimicked by SARS-CoV-2 are MICAL3 (length = 2002 amino acids) and BRI3 (length = 125 amino acids) respectively.

The sequence conservation of each mimicked peptide was derived from all the 46,513 sequenced SARS-CoV-2 genomes available in the GISAID database (as on 06/13/2020).

The immune epitope database (IEDB)^11^ was used to examine the experimentally established, in-vitro evidence for MHC presentation against human or SARS-CoV antigens. The peptides of potential immunologic interest were identified from the IEDB database using the following pair of assays. One of the assays involved purification of specific MHC-class I alleles and estimating the K_d_ values of specific peptide-MHC complexes through competitive radiolabeled peptide binding^12^. The other assay uses mass spectrometry proteomic profiling of the peptide-MHC complexes, where the MHC complexes were purified from the cell lines specifically engineered to produce mono-allelic MHC class I molecules. The identity of the peptide sequence bound to the Class-I MHC molecules was elucidated using mass-spectrometry^13^.

### Analysis of RNA expression in cells and tissues

A distribution of RNA expression across all the expressing samples collected from GTEx, Gene expression omnibus, TCGA and CCLE is created. In this distribution, a high-expression group is defined as the set of samples associated with the top 5% of expression level. Enrichment score captures the significance of the token in the high-expression group. The significance is captured based on Fisher’s test along with Benjamini Hochberg correction. In comparison of gene expression across tissue types in GTEx, the specificity of expression is computing using ‘Cohen’s D’, which is an effect size used to indicate the standardised difference between two means.

### Analysis of overlapping peptides between the proteome and viral proteomes

We analyzed the protein sequences from 9434 viral species (taxons) from ncbi refseq (https://ftp.ncbi.nlm.nih.gov/refseq/release/viral/). On average around 45.57 unique 8mer/9mers per viral taxa were detected to be identical to a known human protein (mean = 45.57, median = 12, sd=±135.38). The highest number of identical peptides were detected in ‘Pandoravirus dulcis’ virus with 4423 peptides having an exact identical match to one or more human proteins.

## Results

### Identifying specific human peptides mimicked by SARS-CoV-2 and with in-vitro evidence for MHC binding

A set of 20 8-mer/9-mer peptides are mimicked by SARS-CoV-2 and no other human coronaviruses (see **Methods**). Of these 20 peptides, four peptides are constituted within established MHC-binding regions (**Table 1 - Panel A**). The 4 peptides with specific MHC-binding potential that are novel to SARS-CoV-2 map onto the following human proteins: Alpha-Amidating Monooxygenase (PAM), Annexin A7 (ANXA7), Peptidylglycine 6-phosphogluconate dehydrogenase (PGD), and Centromere protein I (CENPI) (**Figure 1d**). Analyzing the sequence conservation of the SARS-CoV-2-exclusive peptides shared with the above 4 human MHC binding peptides, shows that these SARS-CoV-2 peptides are largely conserved till date (**Table 2**). The previous human-infecting coronavirus strains (SARS-CoV, MERS, seasonal HCoVs) are notably bereft of these novel SARS-CoV-2 epitopes. An alternatively spliced variant of the human arachidonate 5-lipoxygenase activating protein (ALOX5AP - ENSP00000479870.1; ENST00000617770.4) contains an 8-mer peptide that is mimicked by SARS-CoV-2 as well as SARS-CoV, but not any of the seasonal HCoVs. This peptide has in-vitro evidence for positive MHC Class-I binding. Additionally, there are 4 human helicases (MCM8, DNA2, MOV10L1, ZNFX1), each containing peptides with established evidence of MHC Class-I binding that are also mimicked by SARS-CoV-2 and by previous human-infecting coronaviridae strains (**Table 1 - Panel B; Figure 1d**)(**Table 2**).

**Table 2.**
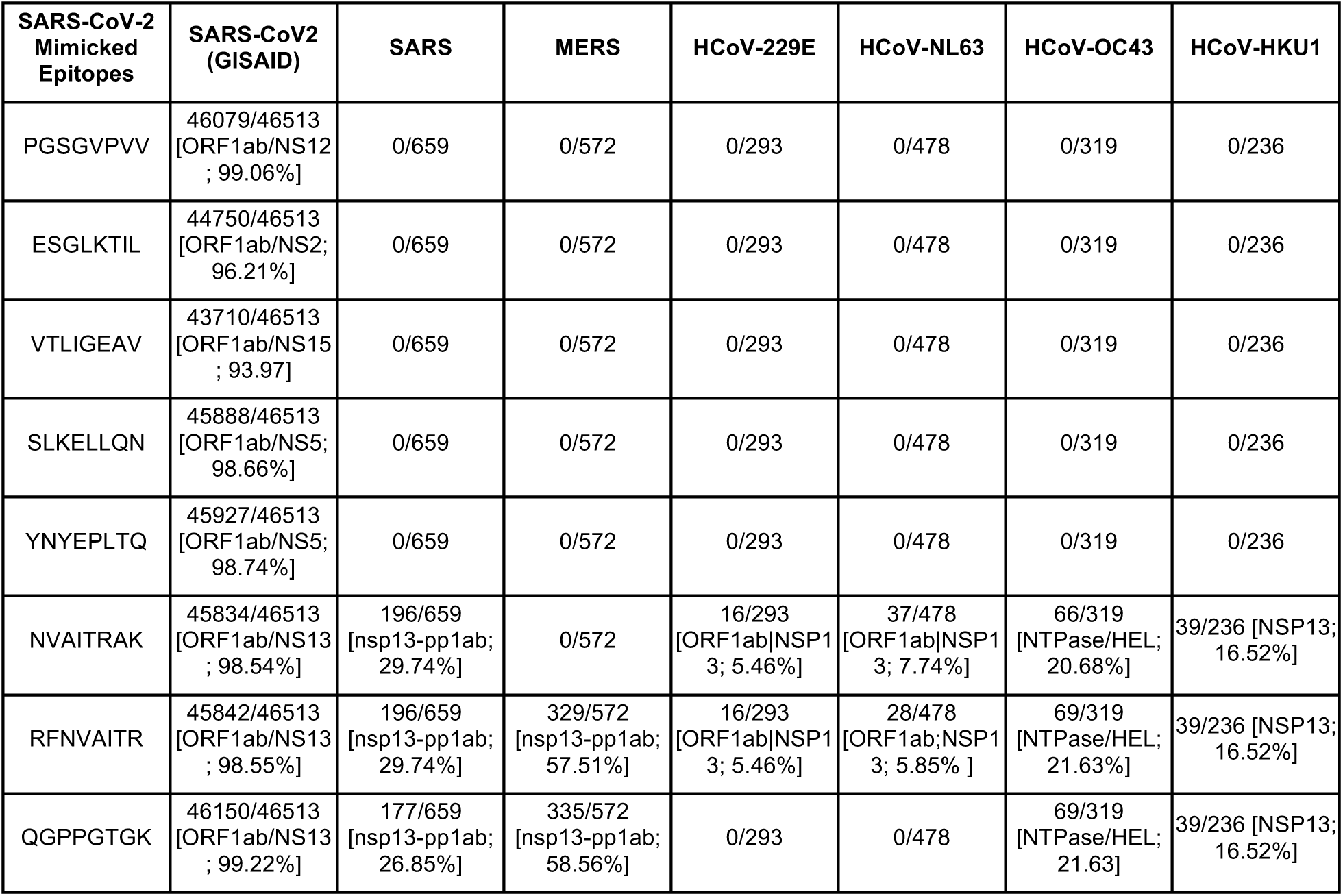
Amino acid sequence conservation of the SARS-CoV-2 peptides mimicking human proteins. The PGSGVPVV peptide from the NSP12 protein is present in 46079 out of 46513 SARS-CoV-2 sequences (99.1% conserved; mimics human PAM protein), the ESGLKTIL peptide from the NSP2 protein is present in 44750 out of 46513 SARS-CoV-2 sequences (96.2% conserved; mimics human ANXA7), the VTLIGEAV peptide from the endoRNAase protein is present in 43710 of 46513 SARS-CoV-2 sequences (94% conserved; mimics human PGD); and the SLKELLQN peptide from the 3C-like proteinase is present in 45888 of 46513 SARS-CoV-2 sequences (98.7% conserved; mimics human CENPI). Furthermore, the PGSGVPVV (NSP12 peptide mimicking PAM), ESGLKTIL (NSP2 peptide mimicking ANXA7), VTLIGEAV (endoRNAase peptide mimicking PGD), and SLKELLQN (3C-like proteinase mimicking CENPI) were not found in any of the proteins from seasonal coronavirus strains downloaded from ViPRdb as on 06/15/2020 -- HCoV-229E (756 protein sequences), HCoV-HKU1 (1310 protein sequences), HCoV-NL63 (1462 protein sequences), and HCoV-OC43 (1921 protein sequences). The YNYEPLTQ peptide from the 3C-like proteinase is present in 45927 out of 46513 SARS-CoV-2 sequences (98.7% conserved; mimics human helicase MCM8 protein), the NVAITRAK peptide from the viral helicase is present in 45834 out of 46513 SARS-CoV-2 sequences (98.5% conserved; mimics human helicase DNA2), the RFNVAITR peptide from the viral helicase is present in 45842 of 46513 SARS-CoV-2 sequences (98.6% conserved; mimics human helicase MOV10L1); and the QGPPGTGK peptide from the viral helicase is present in 46150 of 46513 SARS-CoV-2 sequences (99.2% conserved; mimics human ZNFX1). Moreover, NVAITRAK; RFNVAITR and QGPPGTGK peptides were found in 158/319 (49.5%), 161/319 (50.5%) and 69/319 (21.6%) strains of HCoV-OC43 in the NSP10 (NTPase/HEL) protein. QGPPGTGK peptide was also found in 39/236 seasonal HCoV-HKU1 strains in the NSP13 protein. YNYEPLTQ peptide was not found in any of the seasonal human coronavirus strains.

### Novel mimicry of human PAM, ANXA7, and PGD by SARS-CoV-2, suggests an enrichment of mimicked peptides in lung, esophagus, arteries, heart, pancreas, and macrophages

Considering bulk RNA-seq data from over 125,000 human samples with non-zero PAM expression shows that PAM is highly expressed in pancreatic islets (enrichment score = 276.9 across 6 studies and 552 samples), artery (enrichment score = 244.5 across 3 studies and 343 samples), heart (enrichment score = 243.6 across 19 studies and 442 samples), aorta (enrichment score = 217.1 across 2 studies and 304 samples), embryonic stem cells (enrichment score = 146.7 across 2 studies and 352 samples), and fibroblasts (enrichment score = 107.7 across 28 studies and 215 samples) (**Figure 2a**). Among 54 human tissues from GTEx, PAM is particularly significant within aortic arteries (n = 432, Cohen’s D = 3.1, mean = 348.2 TPM) compared to other human tissues. Moderate specificity in gene expression is also noted for the atrial appendage of the heart (n = 429, Cohen’s D = 2.2, mean = 461.3 TPM) (**Figure 2b**). Further, immunohistochemistry (IHC) data on 45 human tissues ^14^ shows that the PAM protein is detected at high levels in heart muscles, epididymis, and the adrenal gland (**Figure 2b**).

**Figure 2.**
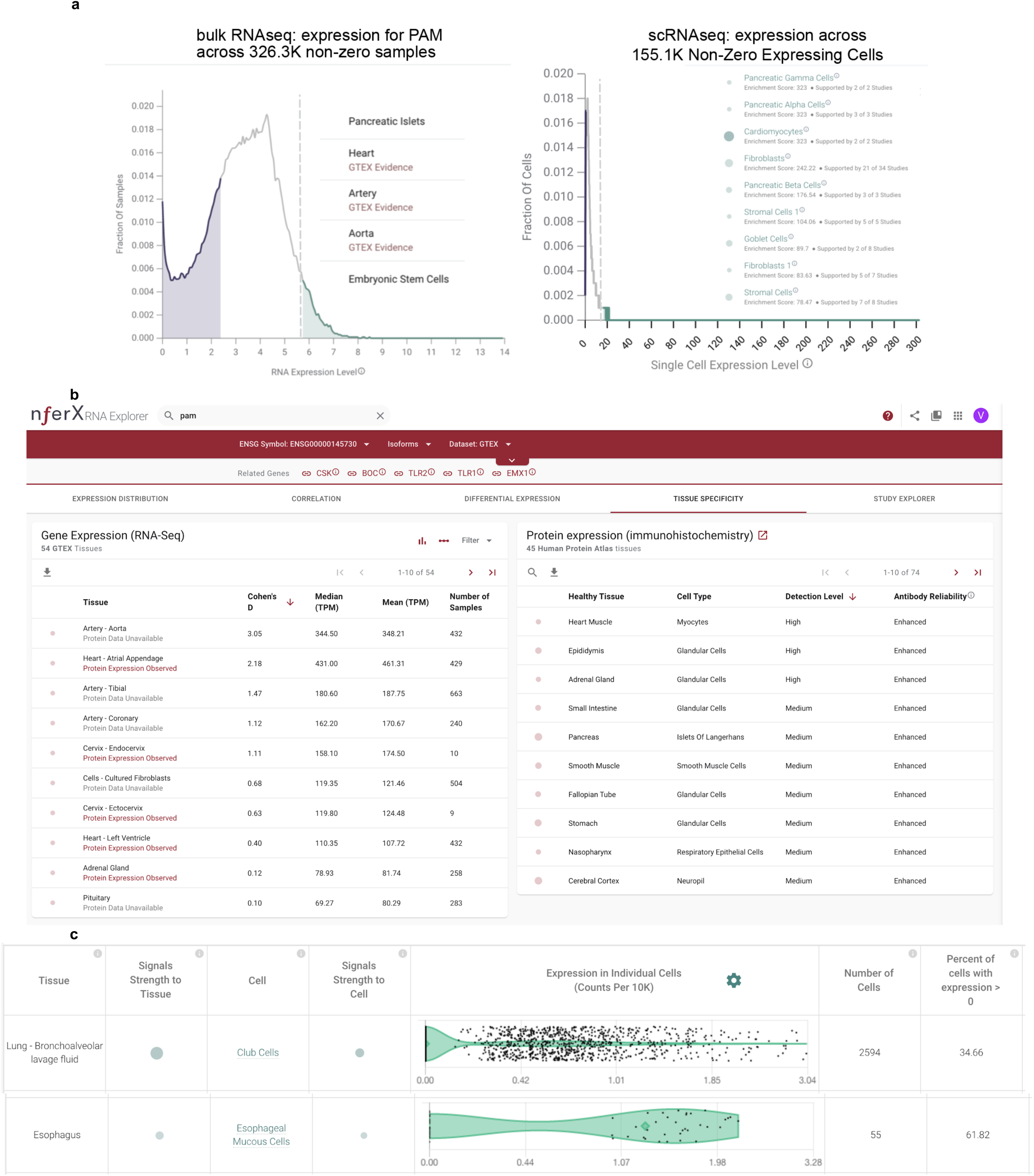
Multi-omics analysis of human PAM. **(a)** (left) Universal bulk RNA-seq analysis of all available human data shows pancreatic islets, heart, artery, aorta and embryonic stem cells harbor PAM significantly. (right) Single cell RNA-seq (scRNA-seq) confirms high PAM-expressing cells include multiple pancreatic cells, cardiomyocytes, goblet cells of the lung, bronchus and intestines, stromal cells of the digestive system, and fibroblasts of multiple organs including the lung, trachea, bronchus, intestines, and heart. **(b)** Analysis of tissue-specific expression pattern of PAM from bulk RNA-seq (GTEx) and triangulation with IHC antibody staining data (HPA) suggests artery, aorta, and myocytes of the heart muscle as significant PAM-expressing tissues. **(c)** Severe COVID-19 patient’s lung bronchoalveolar lavage fluid shows high PAM expression in club cells, which also express the SARS-CoV-2 receptor ACE2 (<u>nferX scRNAseq app - Lung Broncheoalveolar Lavage Fluid</u>).

Exploring all available human Single Cell RNA-seq (scRNA-seq) data shows PAM is expressed in nearly 100% of pancreatic gamma cells, alpha cells, beta cells, delta cells and epsilon cells as well as between 50-90% of activated/quiescent stellate cells, acinar cells, endothelial cells, ductal cells of the pancreas (<u>nferX scRNAseq app - Pancreas</u>). It is also expressed in over 80% of cardiomyocytes, and 40-70% of heart fibroblasts, macrophages, endothelial cells, and smooth muscle cells (<u>nferX scRNAseq app - Heart</u>), as well as in 26-27% of lung pleura fibroblasts, stromal cells, and neutrophils (**Figure 2c**, <u>nferX scRNAseq app - Lung Pleura</u>). Moreover, analyzing the scRNA-seq data from severe COVID-19 patient’s lung bronchoalveolar lavage fluid shows high PAM expression in club cells (**Figure 2d**, <u>nferX scRNAseq app - Lung bronchoalveolar lavage fluid</u>), which intriguingly also express the SARS-CoV-2 receptor ACE2 significantly ^5^. Furthermore, esophagus scRNA-seq analysis shows esophageal mucosal cells and stromal cells as significant PAM expressors (**Figure 2e**, <u>nferX scRNAseq app - Esophagus</u>). Finally, rarer cell types such as pulmonary neuroendocrine cells and goblet cells of the lungs, and some common cell types like lung serous cells and respiratory secretory cells also express PAM significantly.

Similar to the expression profile of PAM, examining 130,400 human samples with non-zero ANXA7 expression shows that ANXA7 is highly expressed in pancreatic islets (enrichment score = 286.65; 543 samples; 3 studies) and artery (enrichment score = 161.68; 184 samples; 3 studies) (**Figure 3 - Supplementary Figure 1**). ANXA7 is significantly expressed in the aortic artery (n = 432, Cohen’s D = 2.1, mean = 163.1 TPM) and the tibial artery (n = 663, Cohen’s D = 2.6, mean = 176.4 TPM) (**Figure 3 - Supplementary Figure 2**). Analysis of the scRNA-seq data on ANXA7 confirms expression in endothelial cells across multiple tissues and organs (<u>nferX Single Cell app - Uterus</u>), and also indicates expression in lung type-2 pneumocytes (<u>nferX Single Cell app - Lung</u>), macrophages, oligodendrocytes (**Figure 3a**). Type-2 pneumocytes are noted to express the SARS-CoV-2 receptor ACE2 from scRNA-seq ^5^. Analyzing the lung bronchoalveolar lavage fluid scRNA-seq data from patients with severe COVID-19 outcomes shows macrophages, lung epithelial cells, T-cells, club cells, proliferating cells, and plasma cells are significant expressors of ANXA7 (**Figure 3b**, <u>nferX scRNAseq app - Lung Bronchoalveolar Lavage Fluid</u>). Additionally from the study of normal lungs, activated dendritic cells and lymphatic vessel cells are noted to express ANXA7 significantly (**Figure 3c**, <u>nferX scRNAseq app - Lungs</u>).

**Figure 3.**
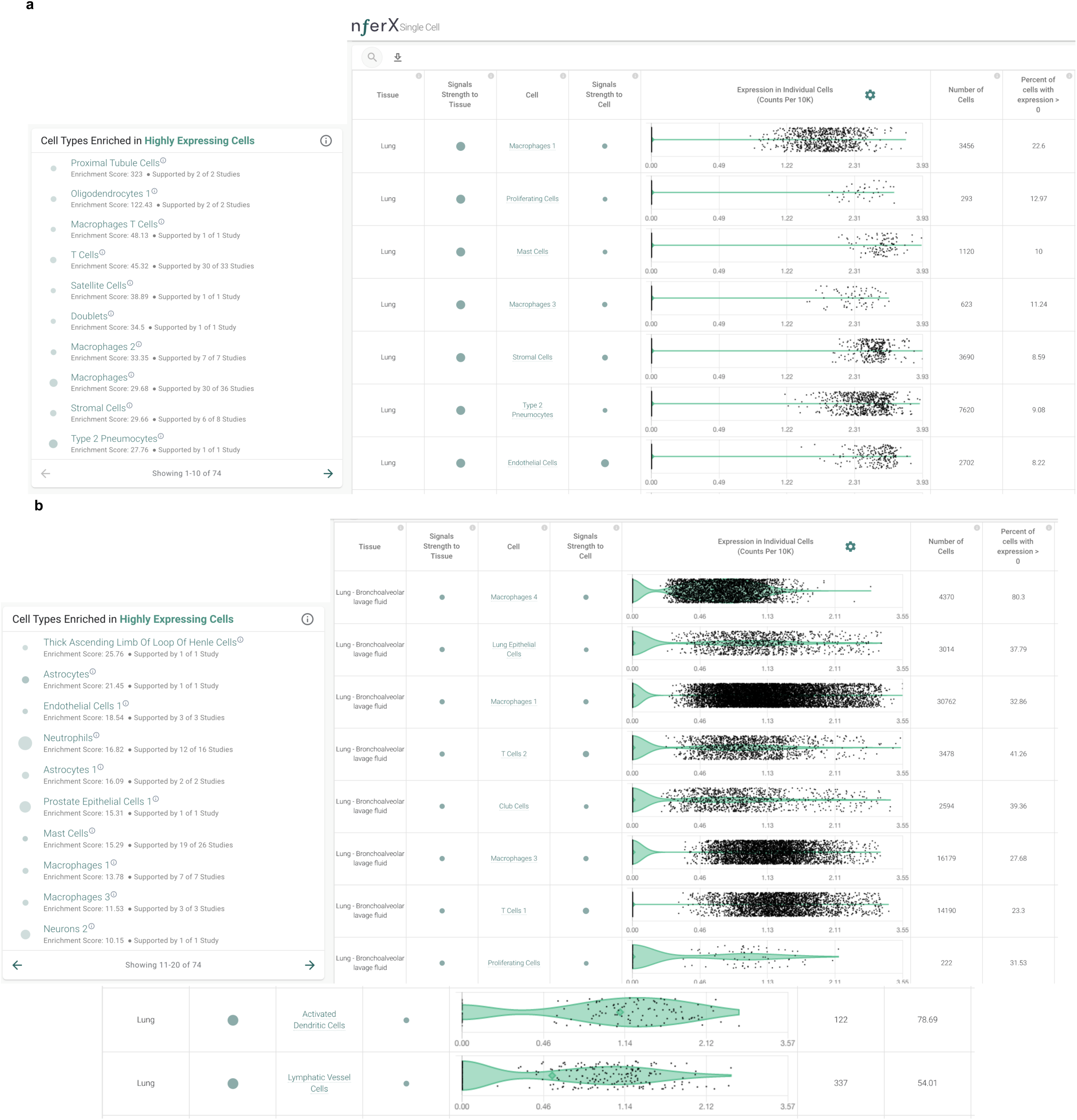
Evidence of ANXA7 and ACE2 expression from human single cell RNA-sequencing data of the lungs. **(a)** List of high ANXA7-expressing cells across human tissues. **(b)** High ANXA7-expressing cells in the lungs include macrophages, proliferating cells, mast cells, stromal cells, Type-2 pneumocytes and endothelial cells; **(c)** Lung bronchoalveolar lavage fluid scRNA-seq shows multiple high ANXA7-expressing cells including macrophages, lung epithelial cells, T-cells, club cells, proliferating cells, and plasma cells (<u>nferX scRNAseq app - Lung Bronchoalveolar Lavage Fluid</u>); **(d)** From the lungs, activated dendritic cells and lymphatic vessel cells are seen to express ANXA7 significantly (<u>nferX scRNAseq app - Lungs</u>).

Assessment of around 128,000 human samples shows that PGD is highly expressed in esophagus mucosa (enrichment = 323, n=510, 2 studies), blood (enrichment = 320.5, n=1020, 39 studies), and macrophages (enrichment = 141.6, n=202, 4 studies). (**Figure 4 - SupplementaryFigure1**). IHC data on 45 human tissues from the Human Protein Atlas ^14^ confirms that PGD is detected at high levels in the esophagus (**Figure 4 - SupplementaryFigure2**), and additionally in the testes, tonsils, bone marrow, gallbladder, spleen and placenta.

Unlike the expression profiles of PAM, ANXA7 and PGD, CENPI’s expression is fairly non-specific and relatively negligible from available data sets. Mild to moderate expression of CENPI is seen in precursor B cells and late erythroid cells, but further studies are needed to ascertain the significance, if any, of CENPI expression, including in the context of COVID-19.

### Multi-pronged mimicry of PAM, ANXA7, and PGD by SARS-CoV-2 and its potential for factoring into the pulmonary-arterial autoinflammation seen in severe COVID-19 patients

Positive HLA-B*40:02 binding has been established for the human PAM peptide (‘KE**PGSGVPVV**L’) and the ANXA7 peptide (‘V**ESGLKTIL**’) ^15–17^, that contain the distinctive mimicking SARS-CoV-2 peptides **(Table 1)**. The closely related HLA-B*40:01 allele also binds this mimicked ANXA7 peptide ^17^. The corresponding mimicking peptides of PAM and ANXA7 are from the viral RNA-dependent RNA polymerase and NSP2 protein respectively, which are constituted within the MHC-binding regions (highlighted above in **bold text**). Additionally, HLA-B*35:01 has experimental evidence for positive binding of the human PGD peptide mimicked by the SARS-CoV-2 virus (**Table 1**).

Given the high expression of ANXA7, PGD and PAM among cells of the respiratory tract, lungs, arteries, cardiovascular system, and pancreas, as well as in macrophages, their striking mimicry by SARS-CoV-2 raises the possibility of individuals with HLA-B*40 and HLA-B*35 alleles being predisposed to potential immune evasion or autoinflammation. Indeed, the potential for broad vascular/endothelial autoinflammation is consistent with the rarer multi-system inflammatory syndrome (MIS-C) or atypical Kawasaki disease noted in few COVID-19-infected children ^18,19^.

### Alternatively spliced human protein variants analysis for mimicry by SARS-CoV-2 highlights another HLA-B*40:01 restricted protein (ALOX5AP) with autoimmune potential

A splicing variant of ALOX5AP (ENSP00000479870.1; ENST00000617770.4) containing the 8-mer peptide ‘PEANMDQE’ is one of 4 alternatively spliced human protein variants that are mimicked by SARS-CoV-2. The other three 8-mer peptides arising from splicing variants do not have any known class I MHC binding reported in the immune epitope database. However, SARS-CoV, which is the only other human-infecting coronavirus in addition to SARS-CoV-2 that also contains **PEANMDQE**, has been experimentally established to possess the **PEANMDQE**SF epitope that positively binds to the HLA-B*40:01 allele.

ALOX5AP is known from literature knowledge synthesis to be associated with ischemic stroke, myocardial infarction, atherosclerosis, cerebral infarction, and coronary artery disease (**Figure 5a**). Single cell RNA-seq studies show numerous types of macrophages expressing ALOX5AP, including in the lungs and brain temporal lobe. Epithelial cells and proliferating cells of the lungs also express ALOX5AP, as do other types of immune cells such as T-cells, neutrophils, and dendritic cells (**Figure 5b**). Taken together with the HLA-B*40:01 restricted binding of the PAM and ANXA7 peptides mimicked by SARS-CoV-2, the ALOX5AP splicing variant also mimicked by SARS-CoV-2 suggests the possibility of immune evasion or autoinflammation in HLA-B*40-constrained COVID-19 patients.

**Figure 4.**
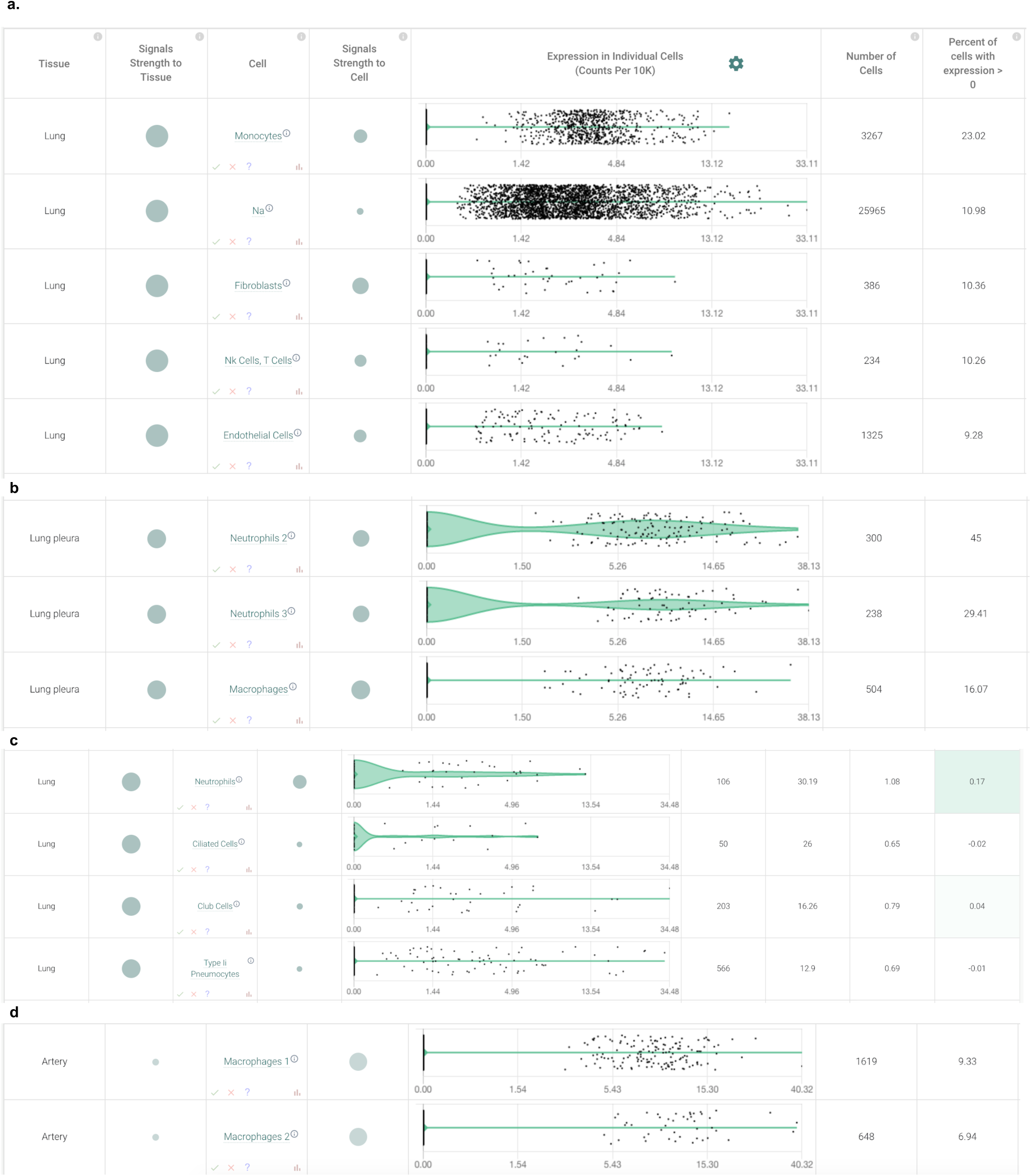
Evidence of PGD expression from human single cell RNA-sequencing data. **(a)** Single cell RNA-seq (scRNA-seq) shows expression in cell types of the human lungs (<u>nferX scRNAseq app - Lungs</u>, <u>nferX scRNAseq app - Lung Pleura</u>, <u>nferX scRNAseq app - Airway Epithelia</u>), and artery (<u>nferX scRNAseq app - Arteries</u>).

**Figure 5.**
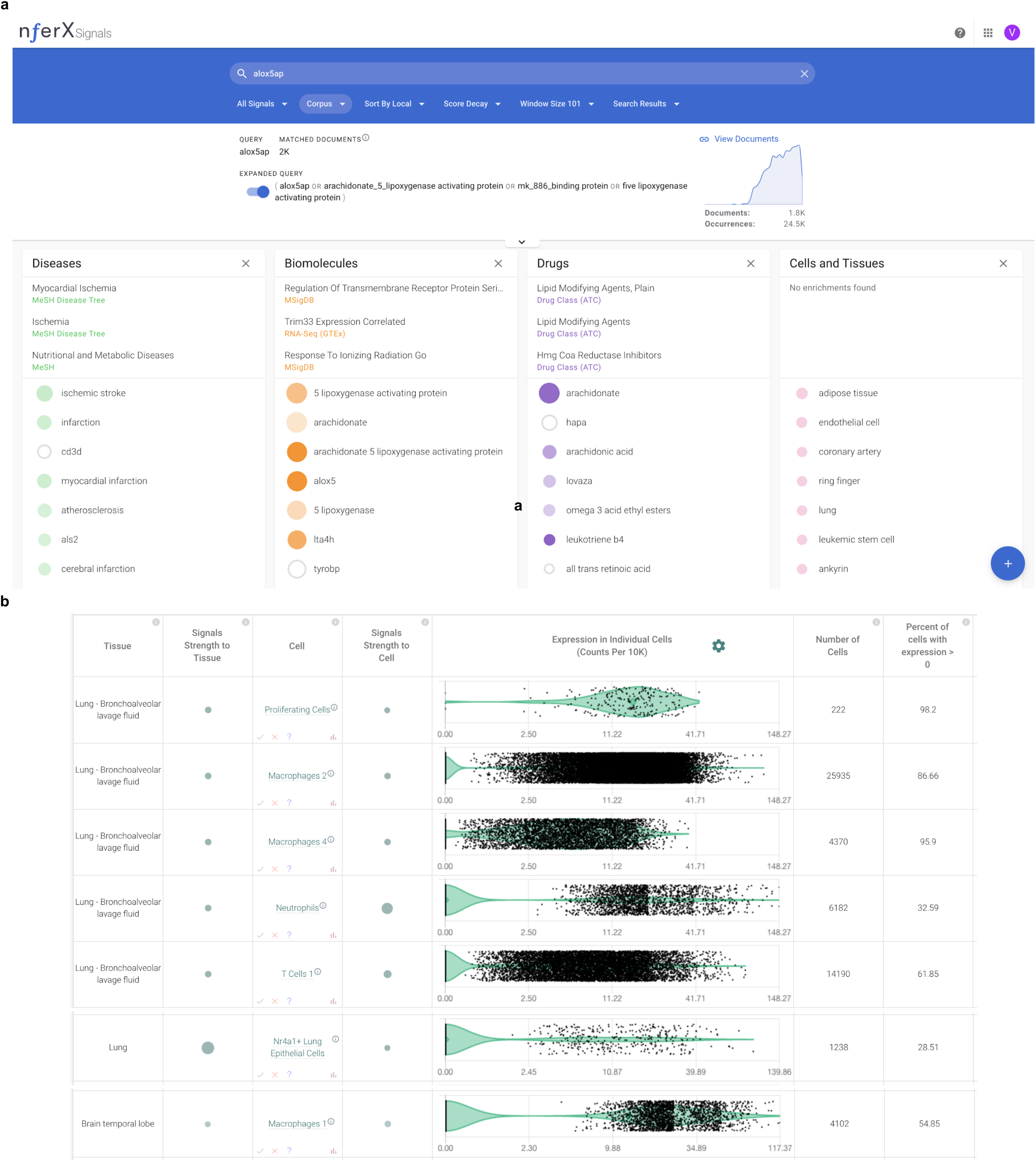
Evidence for ALOX5AP from biomedical knowledge synthesis and single cell RNA-seq. **(a)** Knowledge synthesis suggests involvement of ALOX5AP in ischemic stroke, myocardial infarction, atherosclerosis, cerebral infarction, and coronary artery disease. **(b)** scRNA-seq shows significant expression of ALOX5AP in proliferating cells, macrophages, T-cells, and epithelial cells from the lungs (<u>nferX scRNAseq app - Lung Bronchoalveolar Lavage Fluid</u>, <u>nferX - Lungs</u>) and macrophages of the brain (<u>nferX scRNAseq app - Brain</u>).

### HLA-A*03-binding peptides are shared between helicases of all known human-infecting coronaviridae (HCoVs) and the human proteome

There are seven human protein mimicking peptides that are shared between SARS-CoV-2 and at least one other human infecting coronavirus (**Table 1C**). These proteins include DNA2, MCM8, MOV10L1, ZNFX1, which are all helicases. Analysis of single cell RNAseq suggests that the mimicked human proteins are expressed in neuronal cells and immune cells (**Figure 6c**). The HLA-A*03:01 allele has been established from in-vitro experiments to bind SARS-CoV helicase peptides that mimic an 8-mer peptides from human MOV10L1, DNA2, ZNFX1 and MCM8 helicases (summarized in **Table 1**) ^20^. The HLA-A*31:01 and HLA-A*11:01 alleles, on the other hand, are known to bind peptides containing the human MOV10L1, DNA2, and ZNFX1 helicase mimics; whereas the HLA-A*68:01 allele has established in-vitro evidence of binding peptides containing the human DNA2 and MOV10L1 helicase mimics ^21^. In some of these individuals carrying these HLA alleles (**Table 1C**), a positive T-cell response against their “self” cells that express and display the above coronavirus-mimicked peptides seems plausible.

**Figure 6.**
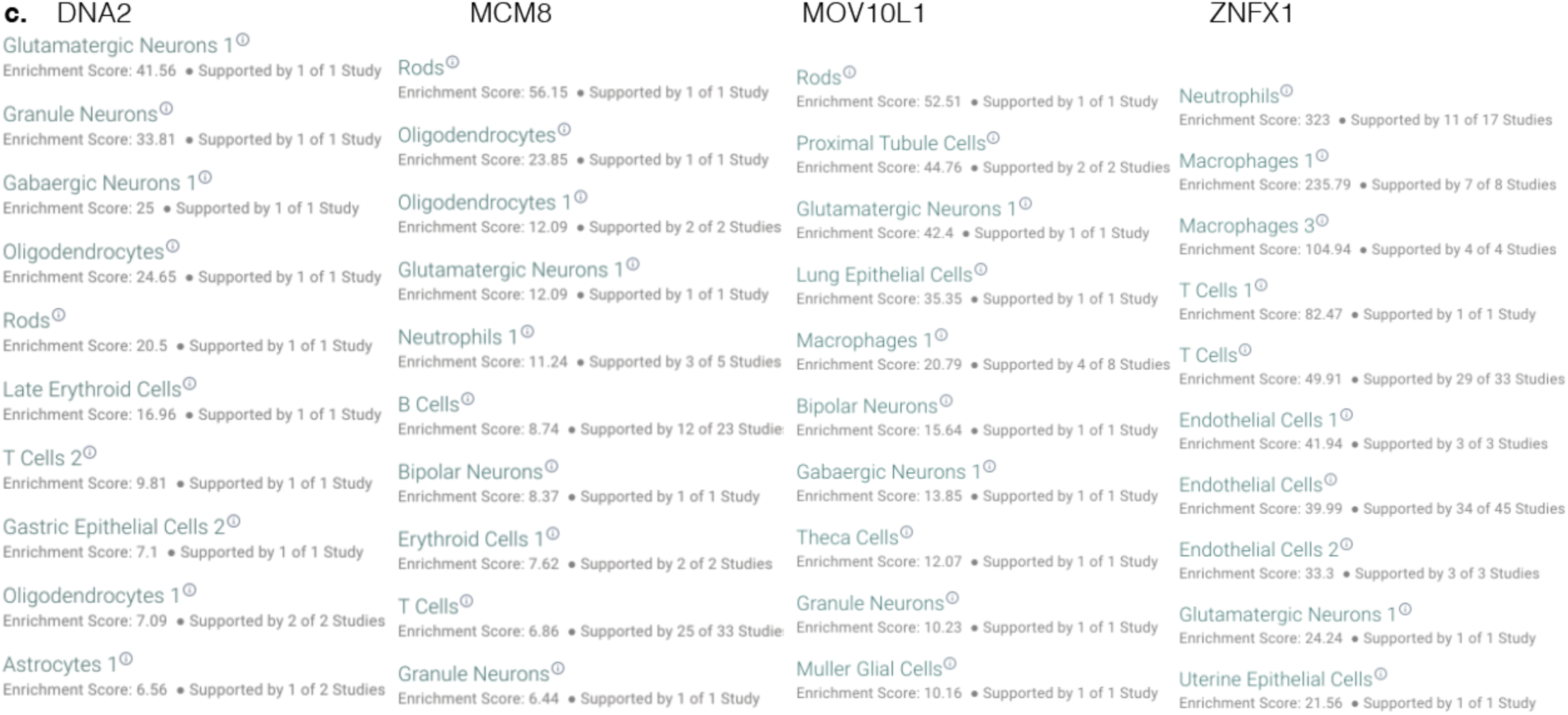
scRNAseq based expression analysis of human helicases containing peptides mimicked by helicases in SARS-CoV-2 and other human coronaviruses.

## Discussion

The presence of identical peptides between viruses and humans has at least two potential implications from an immunologic standpoint. On the one hand, upon presentation of the viral antigens on the surface of infected cells, the virus may evade immune response by masquerading as a host peptide and the recognition of the shared peptides by host regulatory T cells could promote a generally immunosuppressive environment. On the other hand, an autoimmune response can lead to virus-induced autoinflammatory conditions^22^. Either response requires the coupling of both the presence of the appropriate HLA allele and positive T-cell response towards the mimicked peptide epitopes^23^. It is possible that SARS-CoV-2 leverages one or both of these molecular mimicry strategies to exploit the host immune system. In a small minority of patients who happen to have the unfortunate combination of MHC restriction and T-cell receptors as mentioned above, the specific tissues and cell types harboring the mimicked protein would bear the brunt of sustained autoimmune damage. The autoimmune lung and vascular damage reported in severe COVID-19 patient mortality^24–26^ necessitates hypothesis-free examination of both these mimicry strategies.

Our study suggests HLA binding of peptides based on existing literature, but existing literature is by no means exhaustive for identifying HLA binding^11^. There is no known HLA Class-I mediated positive T-cell response against certain 8-mers documented in the immune epitope database. For example, GPPGTGKS peptide is shared by the viral helicase and human VPS4A, VPS4B and SETX. This peptide is also shared with seasonal human coronaviruses (HCoV-OC43; HCoV-HKU1) and previous SARS strains (SARS-CoV; MERS). Further experiments are required to assess any potential for autoinflammation.

Although our current study focussed on human infecting coronaviruses, molecular mimicry is expected to exist beyond human infecting coronaviruses. A stringent BLAST search was also performed for all the four immunomodulatory peptides specific to SARS-CoV-2 against all the sequences of Coronaviridae family in the non-redundant protein database. There were no hits found outside the orthocoronavirinae family for these peptides. An exact match for peptides - ‘PGSGVPVV’, ‘VTLIGEAV’ and ‘SLKELLQN’ was found only in either the pangolin coronavirus or the Bat coronavirus RaTG13. An exact match for ‘PGSGVPVV’ was also found in canada goose coronavirus (YP_009755895.1). The human ANXA7-mimicking peptide ‘ESGLKTIL’ is however noted only in SARS-CoV-2 sequences, with the closest known evolutionary homologs attributed to BAT SARS-like coronavirus (ESGLKTIL), the NL63-related bat coronavirus strains, and the recently sequenced pangolin coronavirus (**Figure 3 - Supplementary Figure2)**.

Our observed multi-pronged human mimicry of SARS-CoV-2, including in peptides that are notably missing from all previously human-infecting coronavirus strains, may in conclusion, owe their origins to zoonotic transmission from coronaviruses circulating within pangolins and bats as natural reservoirs, aided by genetic recombination and purifying selection^27,28^. Our hypothesis-free computational analysis of all available sequencing data, from genomic sequencing and single cell transcriptomics across the host-pathogen continuum, sets the stage for targeted experimental interrogation of immuno-evasive or immuno-stimulatory roles of the mimicked peptides within zoonotic reservoirs and human subjects alike. Such a holistic data sciences-enabled “wet lab” platform for characterizing molecular mimicry and its immunologic implications may help shine a new lens on the relentless evolutionary tinkering that propels the rise and fall of viral pandemics.

## Acknowledgements

The authors thank Patrick Lenehan, Travis Hughes, and Murali Aravamudan for their thoughtful feedback.

## Supplementary Material

**Figure 1 - Supplemental figure 1.**
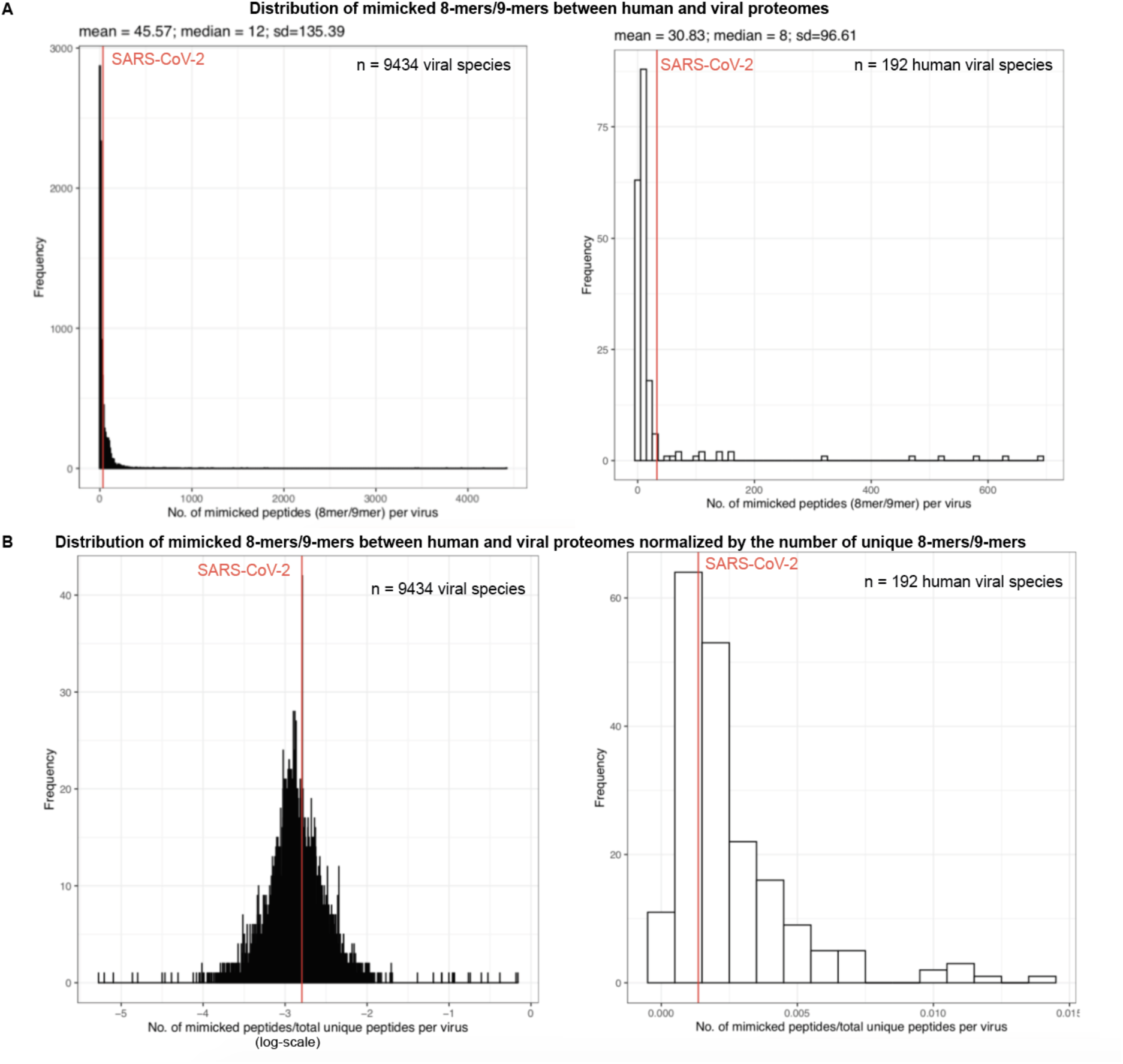
Mimicked 8-mers/9-mers between human and viral proteomes. **(a)** Distribution of mimicked 8-mers/9-mers between human and viral proteomes.**(b)** Distribution of mimicked 8-mers/9-mers between human and viral proteomes normalized by the number of unique 8-mers/9-mers.

**Figure 3 - Supplementary Figure1.**
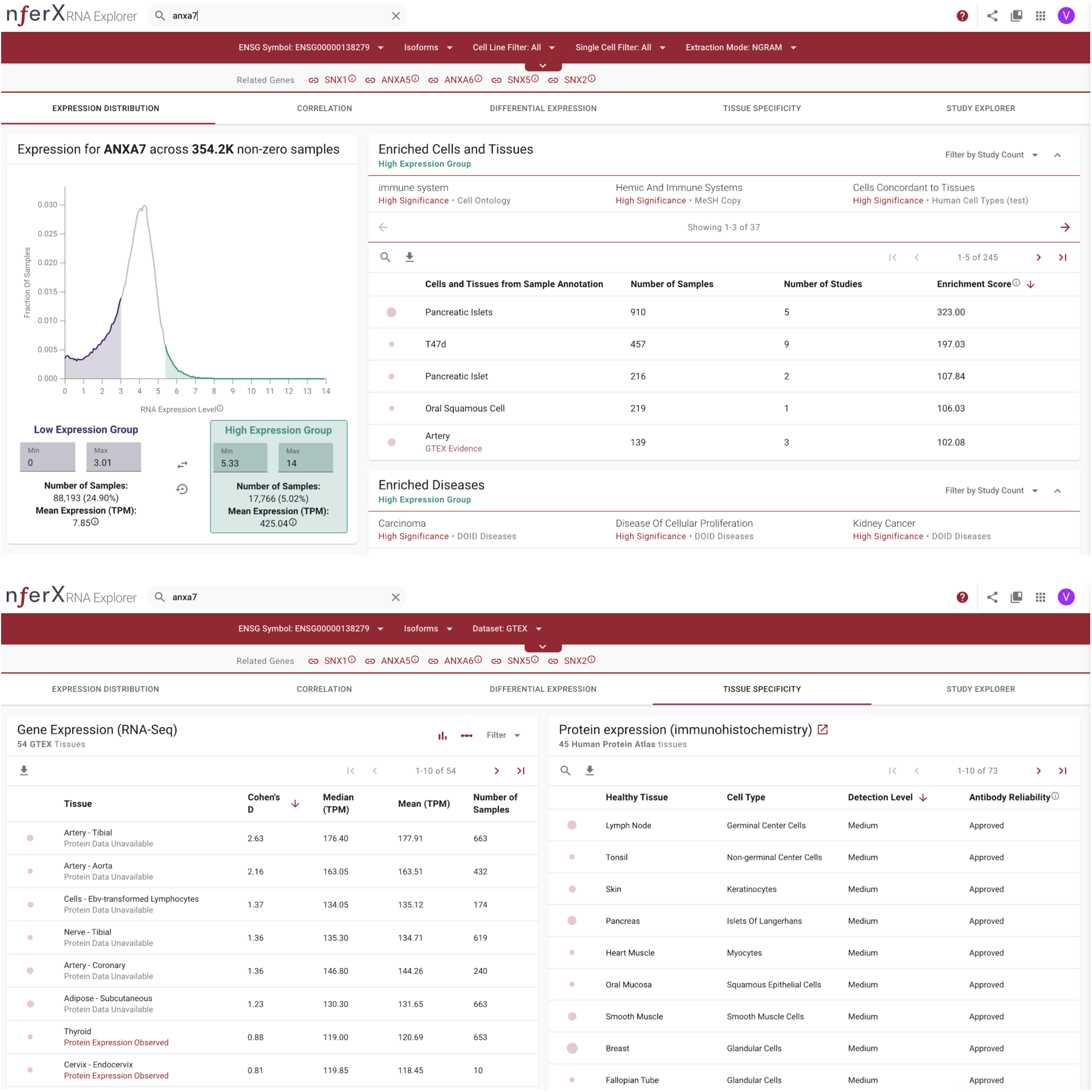
Single cell expression of ANXA7.

**Figure 3 - Supplementary Figure2.**
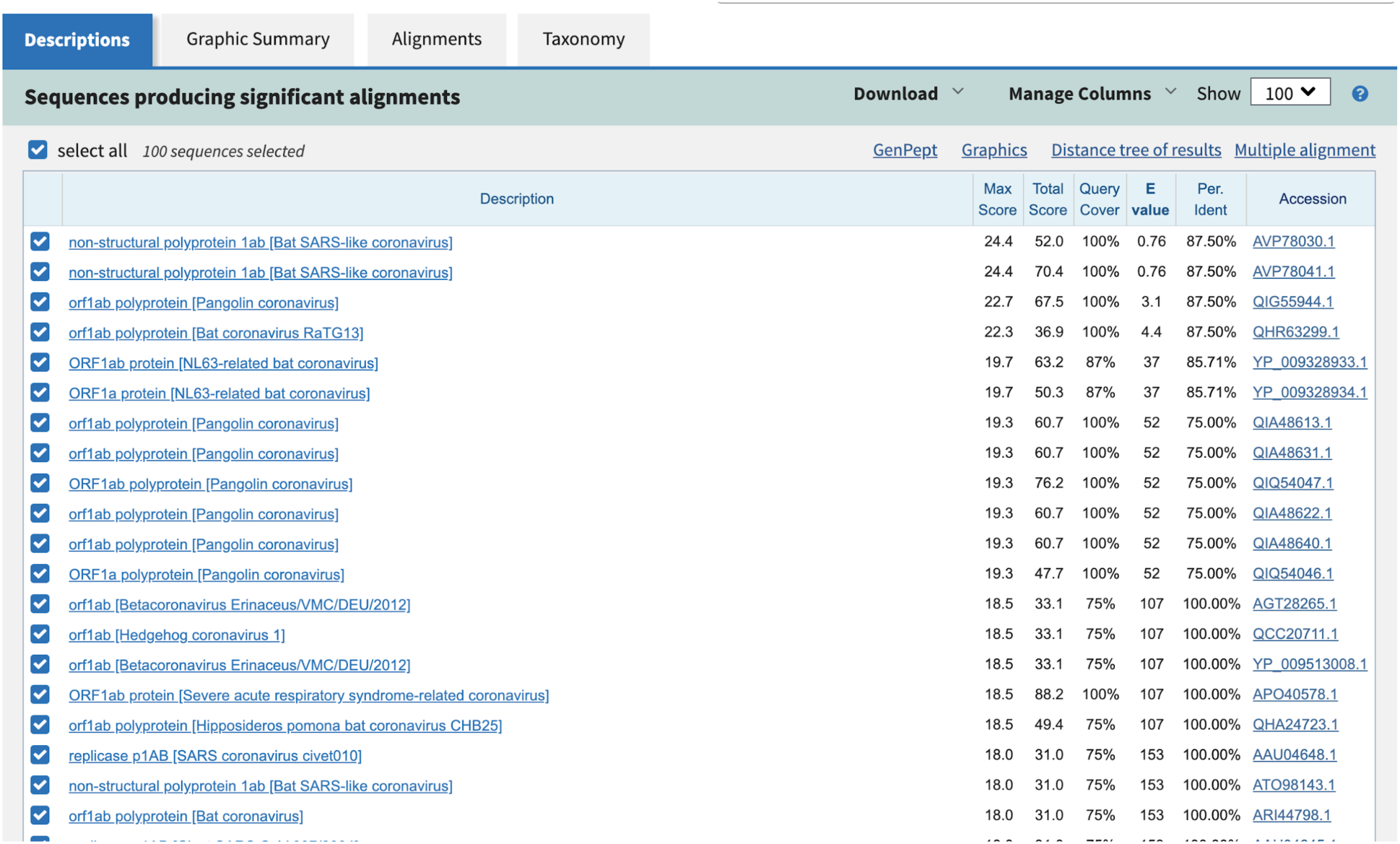
Human ANXA7 mimicking peptide ESGLKTIL is only present in SARS-CoV-2, with the closest known evolutionary homologs being from BAT SARS-like coronavirus (ESGLKTIL), pangolin coronavirus, and the NL63-related bat coronavirus strains.

**Table S1.**
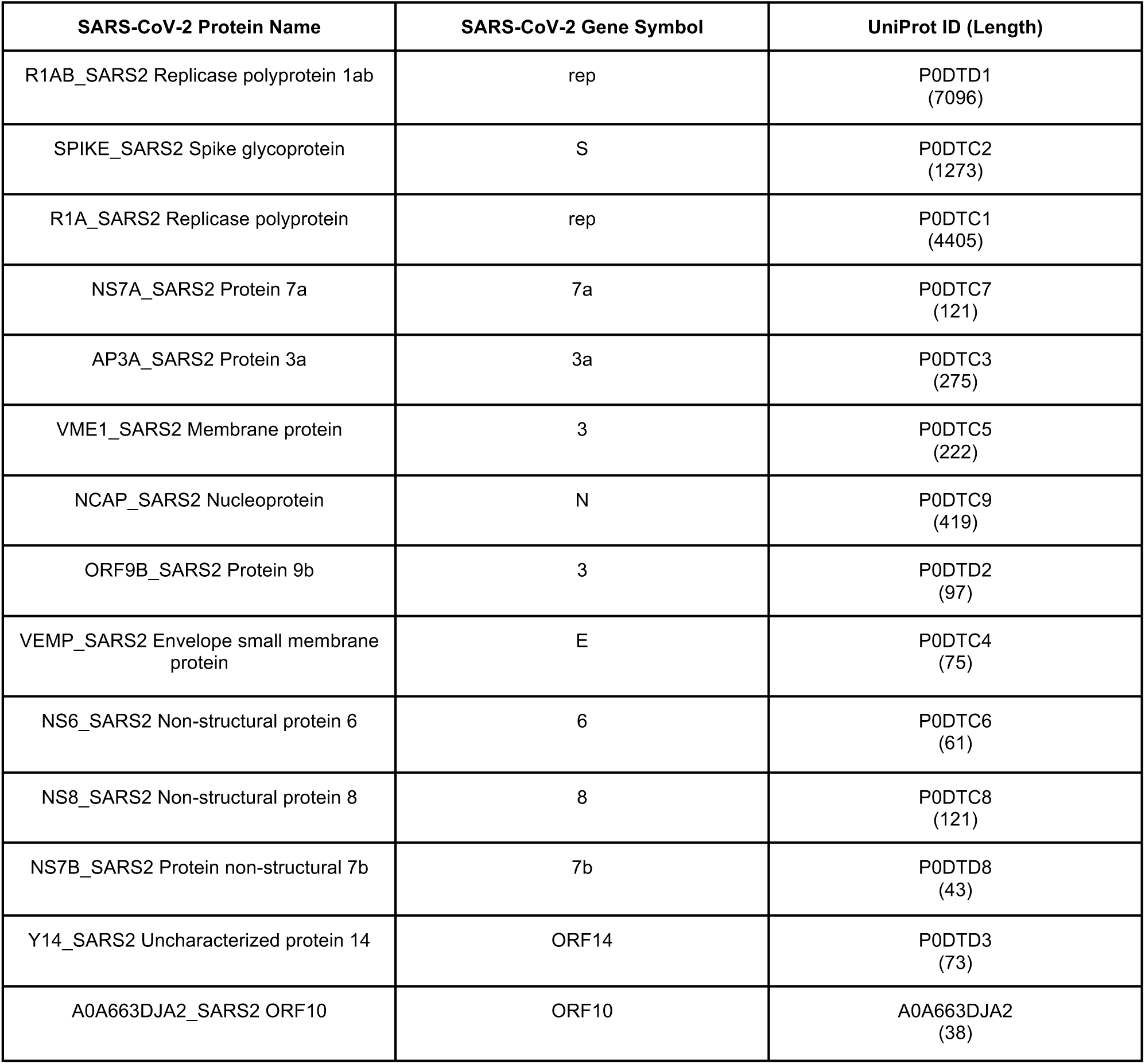
Reference SARS-CoV-2 proteome from UniProt.

**Table S2:**
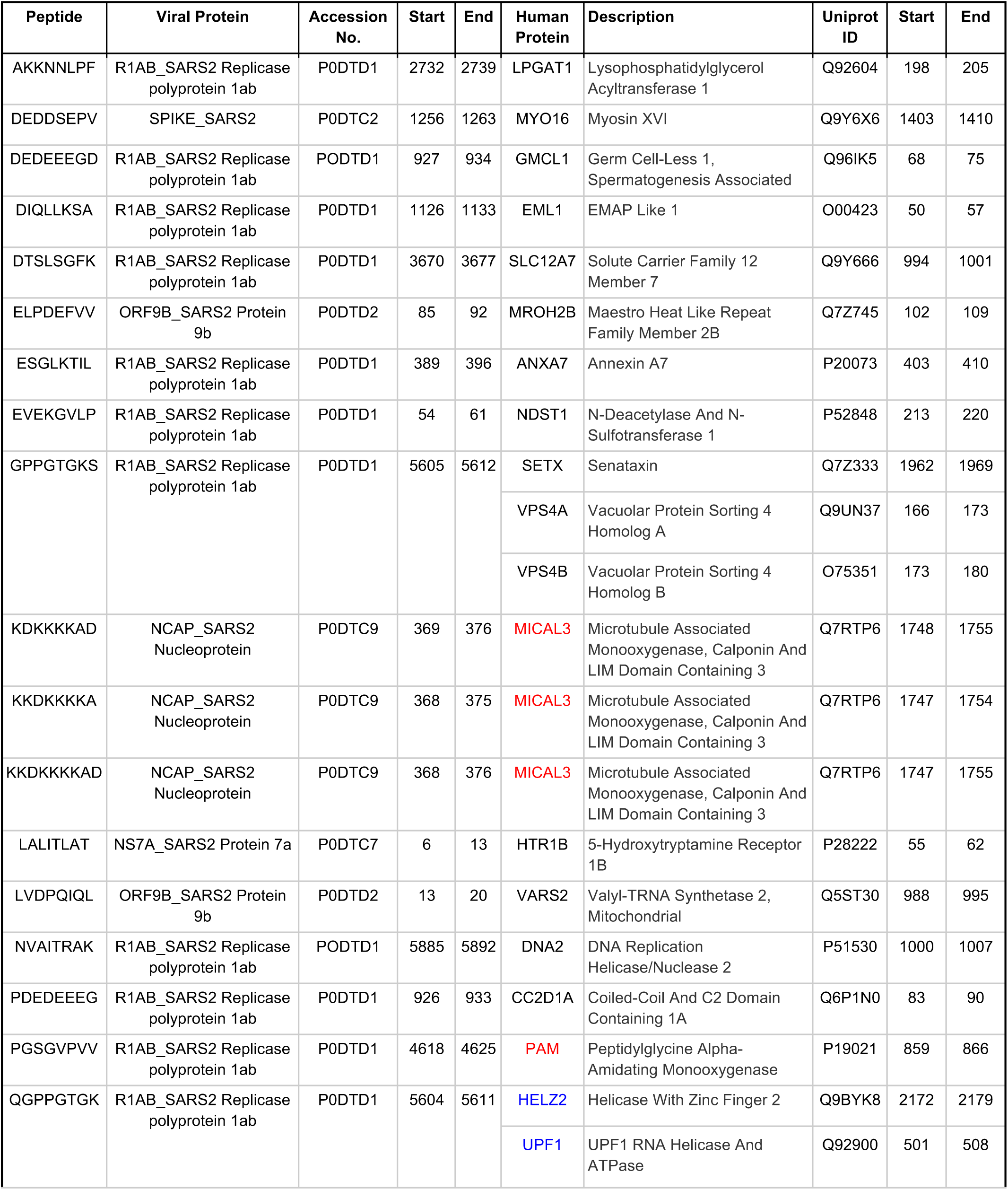
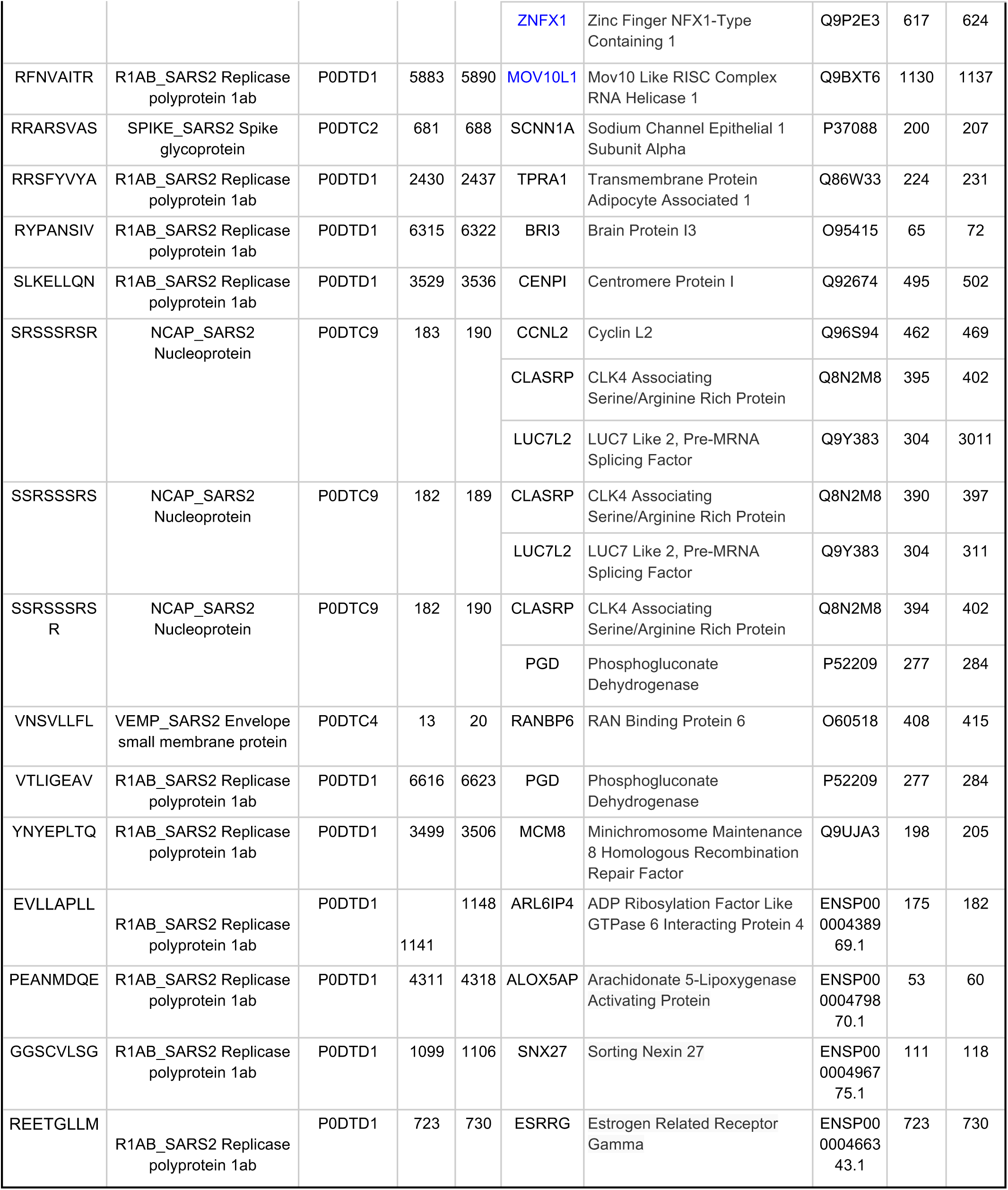
All 33 peptides that are shared between SARS-CoV-2 and the human proteome.

**Table S3.**
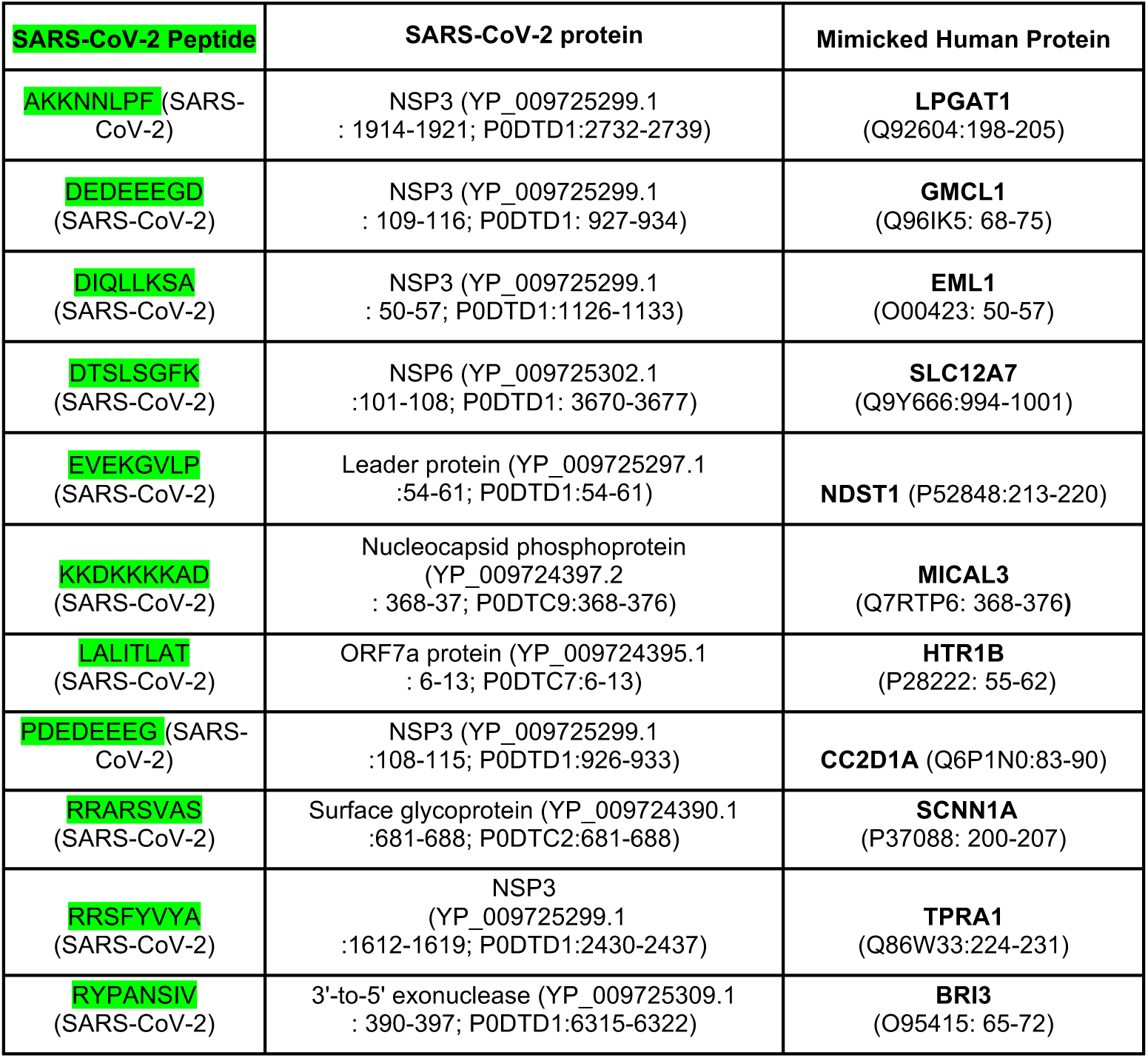
Distinctive peptides from SARS-CoV-2, not present in previously sequenced human coronavirus strains, that do mimic human proteins. There is no compelling positive T-cell immune response against either the human or viral proteins to warrant further discussion in the current study, but these will be the topic of follow-up experimental studies into SARS-CoV-2-based immunologic modulation in humans.

**Table S4.**
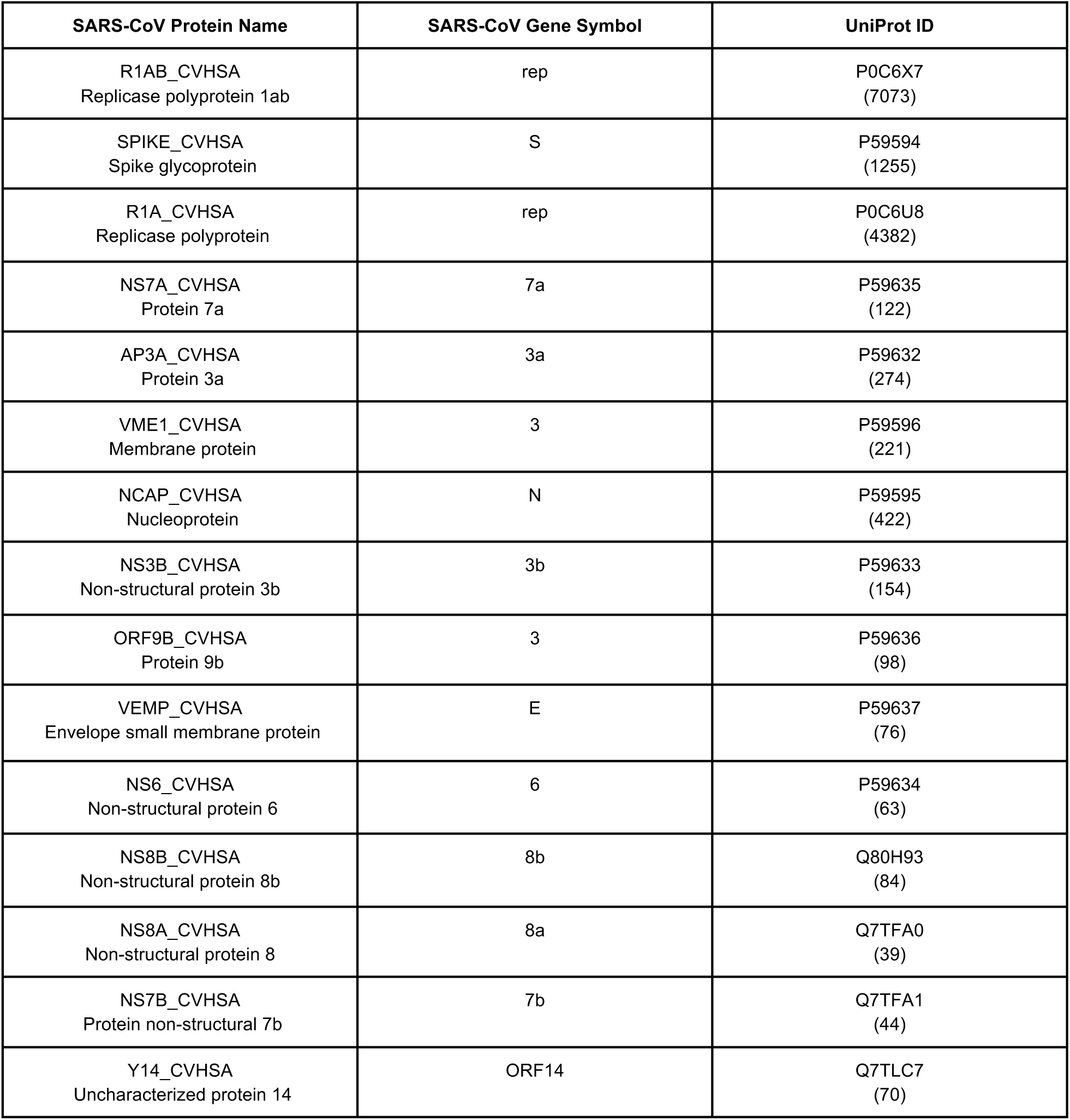
Reference SARS-CoV proteome from UniProt.

**Table S5.**
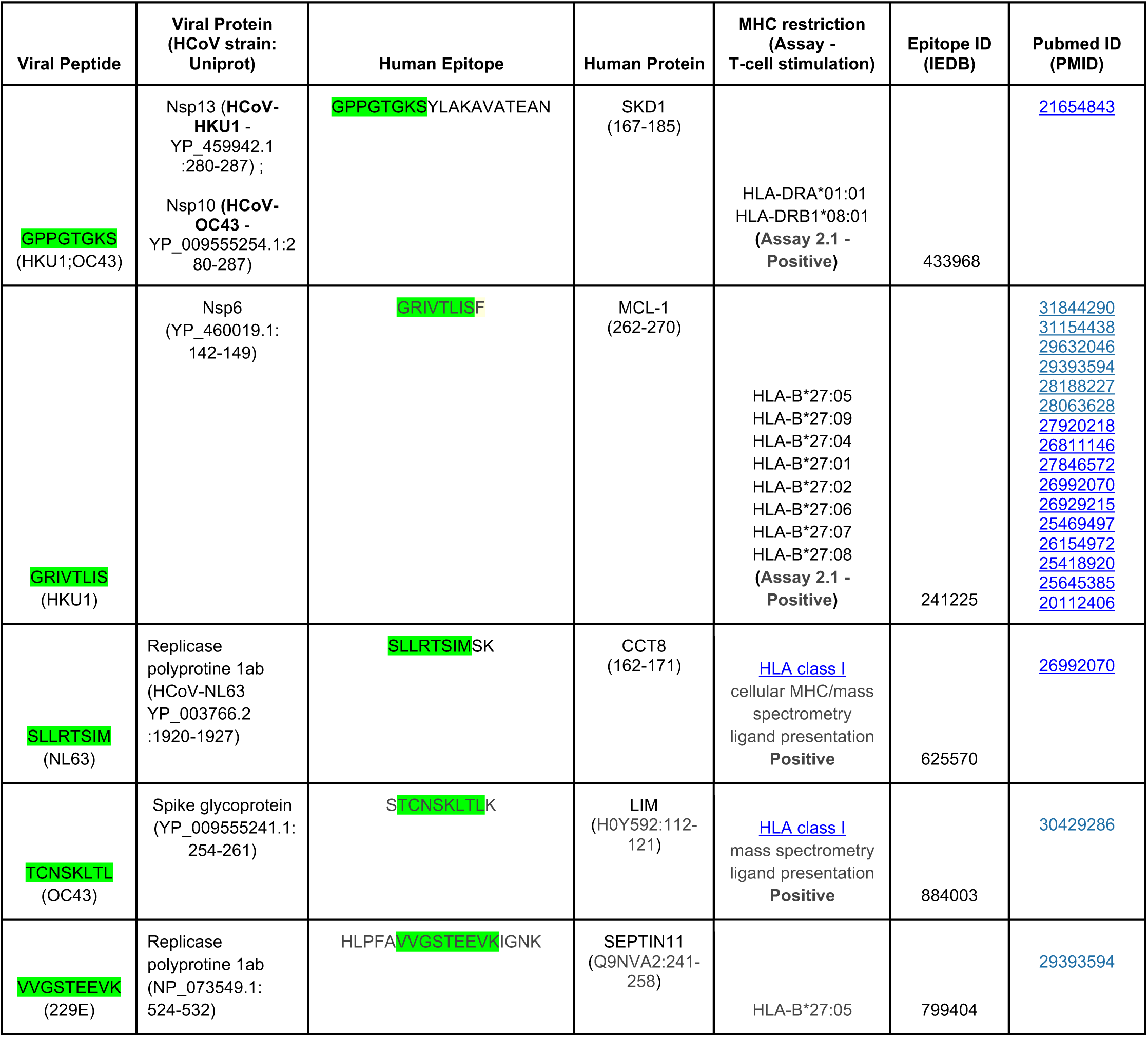
Seasonal human coronavirus (HCoV) peptide mimicry of human proteins with experimental evidence of positive T-cell assays with specific MHC restriction. The MHC-TCR-peptide assays conducted include: (Assay 2.1) Cellular MHC/mass spectrometry, ligand presentation {ref}, (Assay 2.2)

**Table S6.**
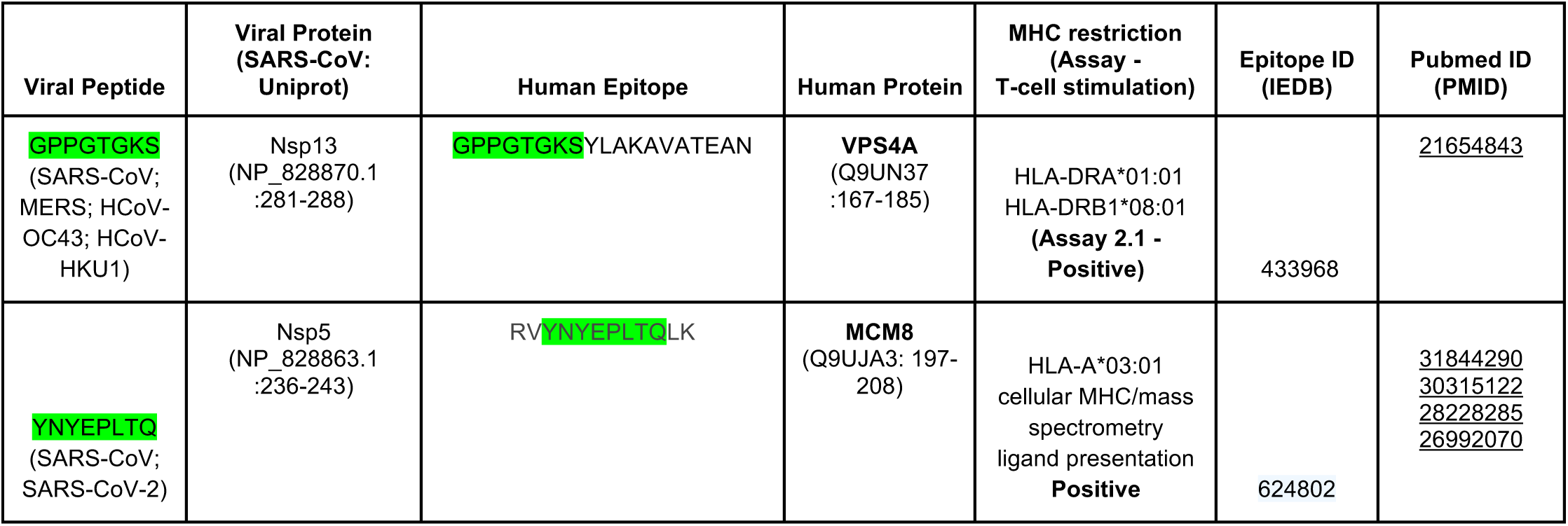
SARS-CoV peptide mimicry of human proteins with experimental evidence of positive T-cell assays with specific MHC restriction. The MHC-TCR-peptide assays conducted include: (Assay 2.1) Cellular MHC/mass spectrometry, ligand presentation {ref}, (Assay 2.2)

**Table S7.**
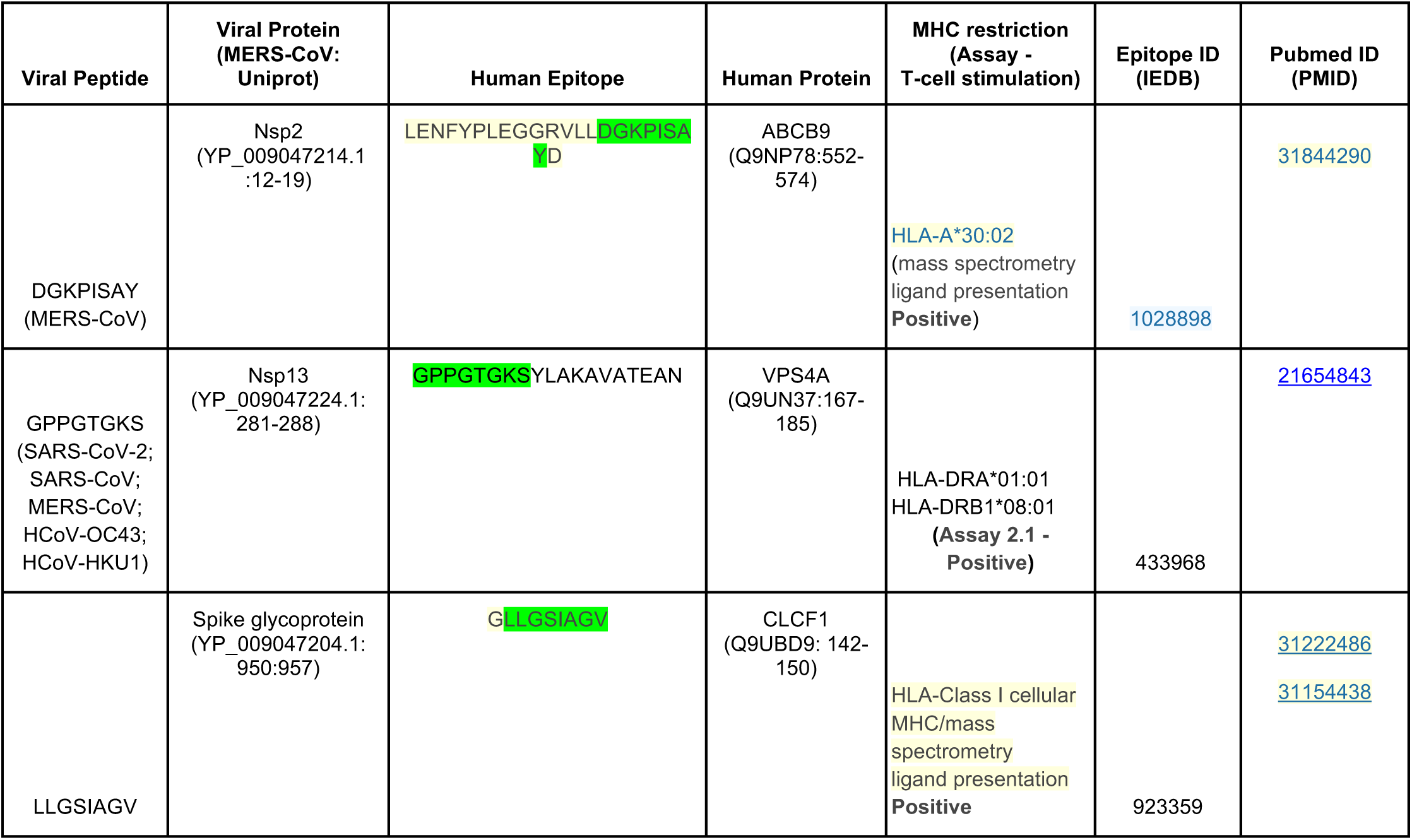
MERS peptide mimicry of human proteins with experimental evidence of positive T-cell assays with specific MHC restriction. The MHC-TCR-peptide assays conducted include: (Assay 2.1) Cellular MHC/mass spectrometry, ligand presentation {ref}, (Assay 2.2)

